# Magnetic Resonance Fingerprinting with Combined Gradient- and Spin-echo Echo-planar Imaging: Simultaneous Estimation of T1, T2 and T2* with integrated-B1 Correction

**DOI:** 10.1101/604546

**Authors:** Mahdi Khajehim, Thomas Christen, J. Jean Chen

**Author notes:** **Corresponding Author**: Mahdi Khajehim, **Email**, **Address**: 3560 Bathurst Street, Toronto, Ontario, M6A 2E1.

## Abstract

**Purpose:** To introduce a novel magnetic-resonance fingerprinting (MRF) framework with single-shot echo-planar imaging (EPI) readout to simultaneously estimate tissue T2, T1 and T2*, and integrate B1 correction.

**Methods:** Spin-echo EPI is combined with gradient-echo EPI to achieve T2 estimation as well as T1 and T2* quantification. In the dictionary matching step, the GE-EPI data segment provides estimates of tissue T1 and T2* with additional B1 information, which are then incorporated into the T2-matching step that uses the SE-EPI data segment. In this way, biases in T2 and T2* estimates do not affect each other.

**Results:** An excellent correspondence was found between our T1, T2, and T2* estimates and results obtained from standard approaches in both phantom and human scans. In the phantom scan, a linear relationship with R^2^>0.96 was found for all parameter estimates. The maximum error in the T2 estimate was found to be below 6%. In the in-vivo scan, similar contrast was noted between MRF and standard approaches, and values found in a small region of interest (ROI) located in the grey matter (GM) were in line with previous measurements (T2_MRF_=88±7ms vs T2_Ref_=89±11ms, T1_MRF_=1153±154ms vs T1_Ref_=1122±52ms, T2*_MRF_=56±4ms vs T2*_Ref_=53±3ms).

**Conclusion:** Adding a spin echo data segment to EPI based MRF allows accurate and robust measurements of T2, T1 and T2* relaxation times. This MRF framework is easier to implement than spiral-based MRF. It doesn’t suffer from undersampling artifacts and seems to require a smaller dictionary size that can fasten the reconstruction process.

## 1) Introduction

Quantitative magnetic resonance imaging (qMRI) typically refers to the quantitative mapping of tissue parameters such as T1, T2 and proton density (PD). Compared to the currently dominant qualitative (e.g. T1 and T2 weighted) techniques, qMRI methods can offer more effective detection and monitoring of different neurological pathologies (1), including stroke (2), neurodegenerative diseases (3–5) and brain tumors (6,7). However, conventional qMRI is limited by very long acquisition times, rendering them unsuitable for routine clinical practice (1,8). Moreover, they suffer from high sensitivity to scanners imperfections (e.g B1 or B0 inhomogeneities). As a result, there is a clear need for quantitative imaging approaches that can estimate multiple tissue parameters in a fast and robust manner.

Magnetic resonance fingerprinting (MRF) is a qMRI method that provides simultaneous estimates of multiple relaxation times as well as information about field-uniformity in a single acquisition (9). In MRF, sequence parameters are varied dynamically and the acquired signal is matched to a pre-calculated dictionary (created using Bloch simulations) using a pattern matching algorithm. The matching identifies the dictionary entry and its corresponding set of predetermined qMRI parameters. So far, MRF has been mostly provided quantification of T1 and T2 relaxation times, and most commonly a spiral readout with a large undersampling factor is used to speed up image acquisition (9–11). Spiral imaging is particularly advantageous for minimizing the echo time (hence off-resonance effects), and has been instrumental in the pioneering work on MRF. The spiral readouts are rotated at each TR to randomize undersampling artifacts (11). Spiral imaging can be used to achieve shorter readout durations than EPI, hence reducing susceptibility effects (12,13). It is also potentially less sensitive to motion (13).

There have been few recent attempts to include T2* in the MRF framework (1,3,14). Wyatt et al. explored a variable-echo-time (TE) scheme to incorporate T2* sensitivity into the originally proposed MRF pulse sequence (15). Hong et al. did so by combining a fast imaging with steady-state precession (FISP) sequence (sensitive to T2) and a multi-echo spoiled gradient echo (SPGR) sequence (sensitive to T2*) (16). Lastly, Wang et al. used a quadratic-RF phase increment pattern to create T2* sensitivity (17). Despite encouraging initial results, these methods exhibit a number of limitations. In Wyatt et al.’s work, the long TEs challenge the design of the undersampled spiral pattern and hence the accuracy of parameter estimation. In Hong et al.’s work, large dictionaries are needed due to the need for simulating numerous effects, which increases the reconstruction times. In Wang et al.’s work, T2 and T2* analyses are coupled so that errors in one estimate can propagate into the second. Furthermore, all of these approaches use undersampled spiral readout with off-line image reconstruction that comes with its own challenges in terms of scan time, ease of implementation and accessibility.

Recently, a non-spiral MRF approach for T1 and T2* quantification has been suggested by Rieger et al. (18,19). A gradient-spoiled gradient-echo (GE) sequence with echo-planar (EPI) readout as well as varying TE, TR, and FA was used for the acquisition. The motivation is that spiral imaging may suffer from image-quality deterioration due to gradient inaccuracies, resulting in the need for custom image reconstruction that is unavailable on commercial scanners. The GE-based implementation by Rieger et al. focuses on T2* instead of T2 estimation.

In this work, capitalizing the flexibility of MRF in terms of sequence design, we extend the approach of Rieger et al. to also quantify T2. We introduce a novel EPI-based MRF approach for simultaneous quantification of T1, T2, and T2* using a combined GE and SE-EPI with integrated B1 correction. We estimate T2 and T2* separately but with the same underlying dictionary. This not only prevents error propagation but also keeps the dictionary size relatively small and easily manageable. Moreover, unlike spiral-based MRF, the raw MRF images have high signal-to-noise (SNR) and are relatively free of artifacts, allowing the use of far fewer frames hence the prospect of faster acquisition. Furthermore, offline reconstruction is not needed, allowing for a more straightforward implementation of MRF.

## 2) Methods

### 2.1) Pulse sequence

A SE-EPI segment was added at the end of the GE-EPI sequence proposed by Rieger et al. (18). A schematic view of the sequence is shown in Figure 1a. This implementation has several advantages. First, as T1 and B1 can both be quantified in the GE-EPI segment and then fed into the dictionary-matching process for the SE-EPI segment, it was found that even as few as 80 volumes (less than half the volumes in the GE segment) were enough for accurate T2 estimation. Second, assuming that T2* can be represented by a monoexponential decay (18), both T2 and T2* can be estimated using the same dictionary, without the need to add another dimension. Third, T2 and T2* estimations are performed from data acquired in separate (GE and SE) halves of the sequence, such that error in one estimate does not affect the other.

**Figure 1.**
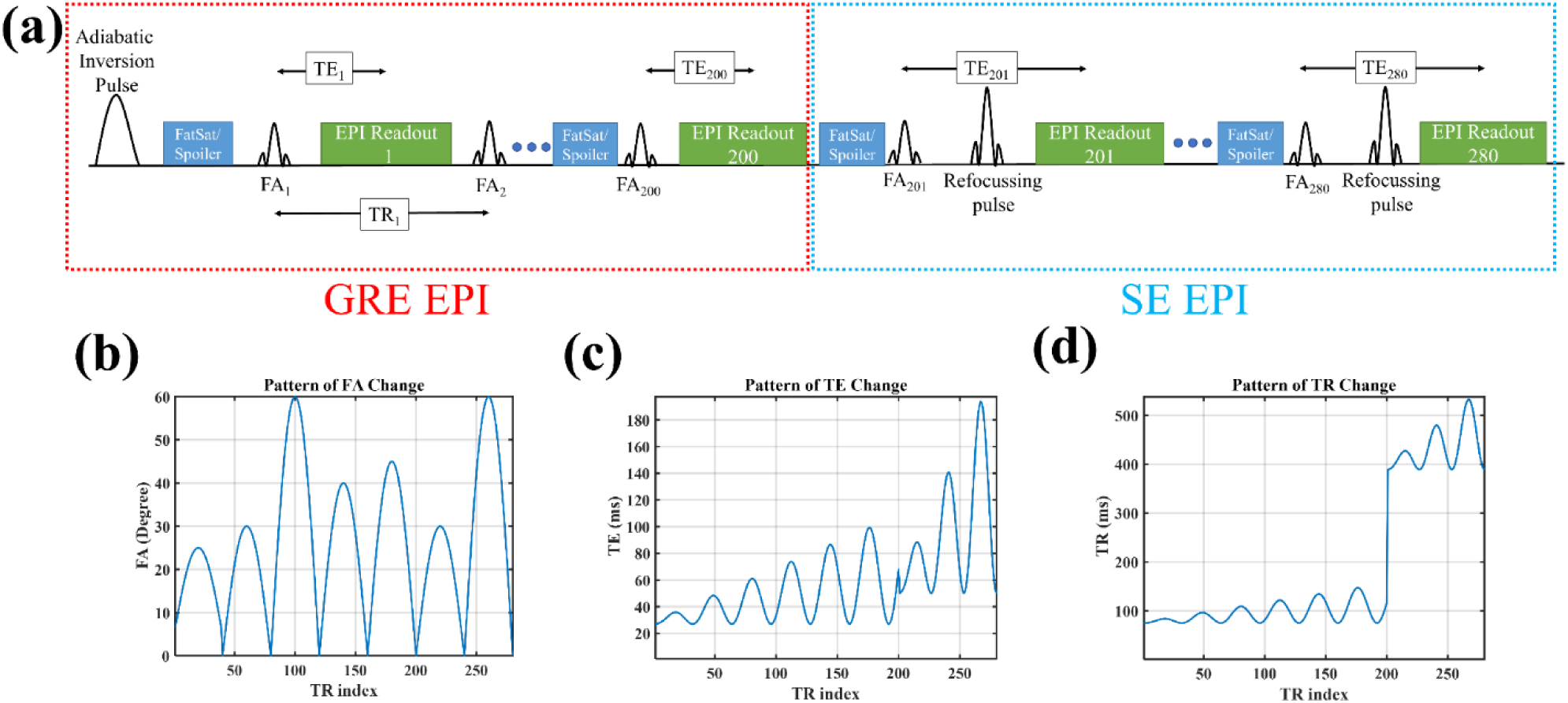
Schematic view of the implemented sequence which is a combination of GE and SE EPI (a). The pattern of FA change in the sequence (b). The pattern of TE change in sequence (c). The pattern of TR change in sequence (d).

To estimate the minimum number of volumes required for the SE-EPI segment, we first obtained a rough estimate of the data SNR (from the GE acquisitions), found to be ∼100. Then, the number of required SE frames was determined through Monte Carlo simulations. We generated a small dictionary with T1 ranging from 500 ms to 1500 ms and T2 ranging from 50 ms to 150 ms (∼1200 entries) while assuming homogeneous B1. Subsequently, Gaussian white noise (based on the estimated experimental SNR) was added to the dictionary. Pattern matching was done (see next section) and the mean relative error of T2 estimation was calculated to investigate the relationship between T2 error and the number of acquired SE volumes. Based on this, we found that 80 SE frames leads to less than 2% error in T2 estimation and seems a reasonable choice by balancing acquisition time and accuracy.

The GE-EPI segment is similar to Rieger et al., with slight modifications in terms of the number of acquired frames and pattern of FA/TR/TE change. After a hyperbolic secant adiabatic inversion pulse, the GE-EPI portion begins: (1) the FA varies in accordance with five half periods of a sinusoidal variation, with FAs ranging overall from 0 to 60 degrees; (2) TEs vary between 25-100 ms while TR is the shortest possible for each TE (range 65-140ms); (3) in addition to fat saturation, both gradient and RF spoiling are implemented using crusher gradients before the fat saturation module (in x, y and z directions). After 200 GE frames, the sequence transitions into SE-EPI for 80 frames --- a slice-selective 180° refocusing pulse is applied before the EPI readout. Crusher gradients are added before and after the refocusing pulse in all three directions to spoil the free induction decay signal that may originate from non-ideal refocusing. In the SE-EPI segment, two half-periods of a sinusoidal function are used for FA variation (with the sinusoidal maxima of 30 and 60 degrees, respectively). High flip angles are avoided to suppress the effect of slice-profile imperfections (20,21). The TE range is 50-190 ms. To counteract the partial saturation of the longitudinal magnetization due to the refocusing pulse and thus to ensure a sufficient level of longitudinal magnetization, a 300 ms recovery time is added after each EPI readout in the SE segment (see Discussion for details). Other sequence parameters common to both GE and SE segments are: matrix size= 128×128, FOV= 220 × 220 mm, voxel size = 1.7×1.7×5 mm, GRAPPA factor=2, #reference lines=62, no partial Fourier, BW/Pixel=1562 Hz, Total acquisition time per slice is ∼45 s. In this proof-of-concept study, single-slice acquisition was used.

### 2.2) Dictionary generation and pattern matching

As was done by Rieger et al. (18), each voxel is represented by one isochromat. The dictionary is generated using the discrete form of the Bloch equations. RF pulses are assumed to be instantaneous, and the RF-slice profile is ignored for simplicity. B1 inhomogeneity is explicitly included into the model as a scaling factor applied to the nominal FAs. Considering min:step:max to represent the range of simulation values, the T1 range was 50:25:2500 ms, the T2/T2* range was 5:5:250 ms and relative B1 was 0.5:0.05:1.5. Overall the dictionary has ∼ 100,000 entries and requires 200 MB of storage. Dictionary generation takes less than 10 minutes on a desktop computer running on a quad-core 1.6 GHz CPU. A two-step dictionary-matching algorithm was implemented. Notably, we assume that the transverse signal decay in the GE segment is characterized by T2* relaxation time while the transverse signal decay in the SE segment is characterized by T2 relaxation time. In the first step, the GE data are used to obtain estimates of T1, T2* and B1, whereas in the second step, the SE data are used to only match for T2, after adopting the T1 and B1 values derived from the GE data. Pattern matching is done using the magnitude of the MR signal and the pre-calculated dictionary using a maximum dot product approach. Dictionary generation and data matching both were implemented in MATLAB R2018b (MathWorks, Natick, MA, USA) using custom-built codes.

### 2.3) Phantom validation

All imaging was performed on a Siemens TIM Trio 3 T system (Erlangen, Germany) using a 32-channel head coil. For validation, phantoms were built with varying concentration of copper (II) sulfate (CuSO_4_) and agar (1-4% w/w). Seven small vials with different T1/T2/T2* properties were placed inside a larger cylindrical plastic container filled with CuSO_4_ doped distilled water. Our MRF estimates were compared to those obtained from standard relaxometry techniques. To validate T1 estimations, inversion recovery turbo SE (IR-TSE) was used with eight different inversion times (TIs=25, 50, 100, 200, 400, 800, 1600, 3200 ms), TR=10s, TE=9.5 ms, BW/Pixel=200 Hz and turbo-factor=8. For T2 validation, a single echo TSE sequence was repeated with seven echo times (TEs=19, 38, 57, 76, 95, 114,132,152 ms), TR=5s, BW/Pixel=200 Hz, Turbo-factor=8. T2* was measured using a single echo GRE repeated for seven different echo times (TEs=5, 15, 30, 45, 60, 75, 90 ms) TR=1s, BW/Pixel=390 Hz and FA=15°. B1 maps were acquired using the double angle method with FLASH sequence, with FA1=60° and FA2=120°, TR/TE=5000/10 ms (22). Standard T1 fitting was done using complex data and the five-parameter model described by Barral et al. (23). A two-parameter model (S=M_0_ exp(TE/T2(*))) was used for both T2 and T2* by fitting a monoexponential curve to the data using nonlinear (Levenberg-Marquardt) curve fitting algorithm (MATLAB). ROIs were manually drawn around each of the small vials with the exclusion of edge voxels. The mean and standard deviation of all the voxels in these ROIs were used for comparisons.

### 2.4) In vivo validation

In-vivo data from a healthy human subject (male, 24 years old) were acquired to validate the sequence, following informed written consent, in accordance to the Institutional Research Ethics Board policy. The sequences and scanner information are similar to those described under “Phantom Validations” and “Pulse Sequence”, except that the standard T2* measurements were obtained using a multi echo-GRE (ME-GRE) sequence with twelve TEs=2-80ms and TR=1000 ms in order to reduce the acquisition time. Manual ROIs (∼ 30 voxels each) were drawn in grey-(GM) and white-matter (WM) regions for subsequent comparisons.

## 3) Results

As a demonstration of image quality, raw MRF images from phantom scans are shown in Figure 2a. Compared to undersampled spiral MRF images (11), our images have high SNR and minimal levels of artifacts. The quality of single-voxel match between dictionary entry and acquired data from the phantom study is illustrated in Figure 2b. Excellent correspondence between the MRF data, fitted dictionary element, and the ground truth dictionary entry (created from standard acquisitions) can be seen. Figure 2c shows an example of the raw MRF images for the human scan. Again, the image is nearly free of artifacts. The correspondence between data, dictionary and ground truth is also satisfactory in this case. As shown in Figure 2d, the in-vivo time-series seems noisier than the phantom data, which is expected due to factors such as motion, physiological noise, or intra-voxel tissue heterogeneity, that are not accounted for in the dictionary generation step. The baseline signal level from the SE segment is lower than from GE section, stemming from the lower baseline longitudinal magnetization. Nonetheless, high accuracy is still achieved since this segment is only used for T2 estimation.

**Figure 2.**
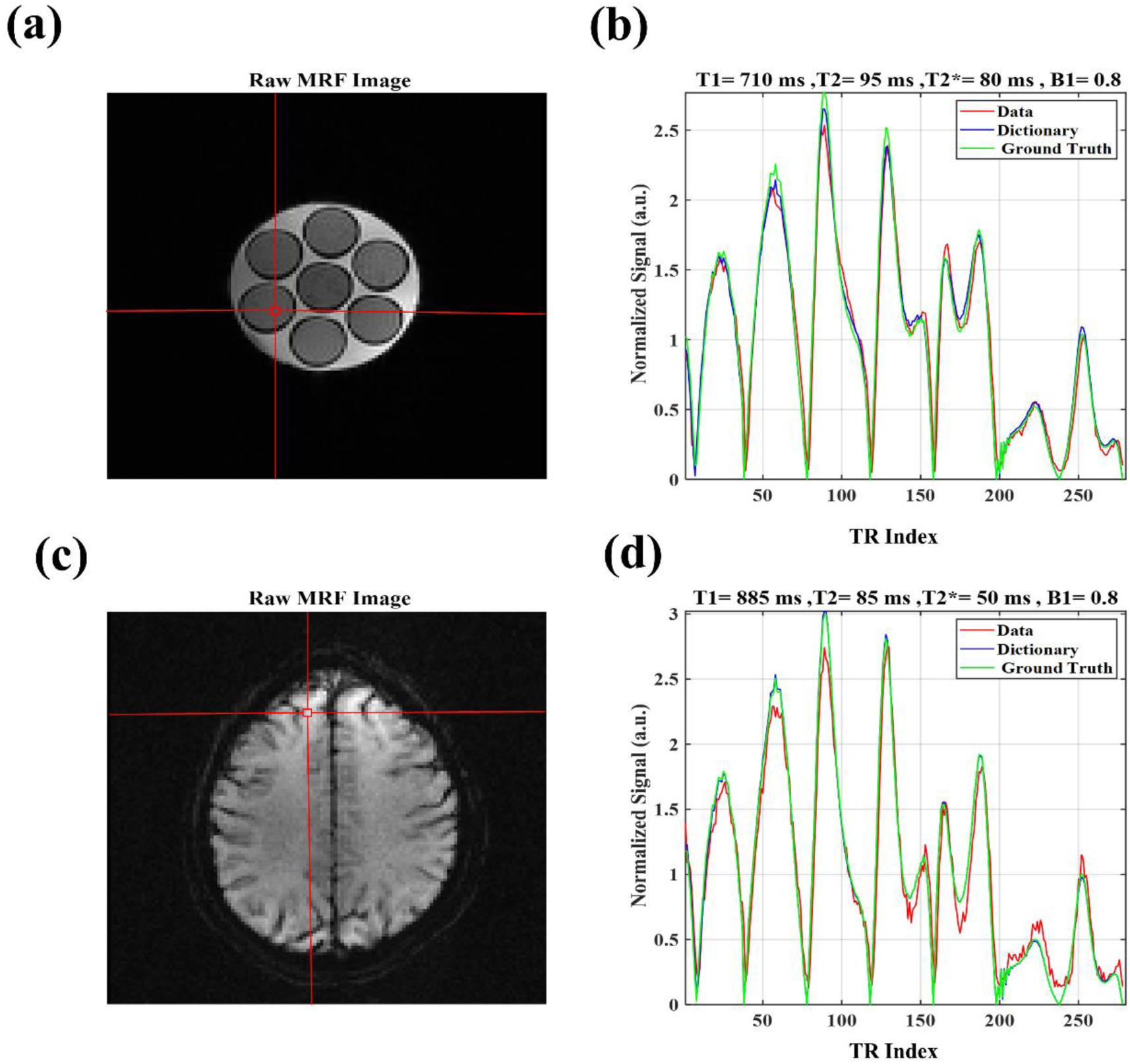
An example of the quality of raw MRF images for the phantom scan (a) along with the acquired time series in red, fitted dictionary entry in blue and ground truth dictionary element in green for one representative voxel (b). Quality of raw image for the human scan (c) along with the correspondence between data, fitted dictionary entry and ground truth dictionary element for one exemplary voxel (d). Ground truth dictionary entry has been created by using relaxometry values that come from the ground truth measurements rather than the MRF scan.

Figure 3 shows the parametric maps obtained from phantom acquisitions using our proposed MRF sequence as well as the standard measurements (ground truth). Excellent correspondence between estimated and expected values (high R^2^ values) are achieved for all three relaxation parameters (T1/T2/T2*). Particularly for T2, the novel contribution of our sequence, the R^2^ is 0.998 and the maximum error is ∼6%. Figure 4 shows the in-vivo MRF results and their correspondence with the standard measurements in terms of image contrast and the range of estimated values. Slight contrast differences can be seen in the CSF and could be due to motion and flow effects.

**Figure 3.**
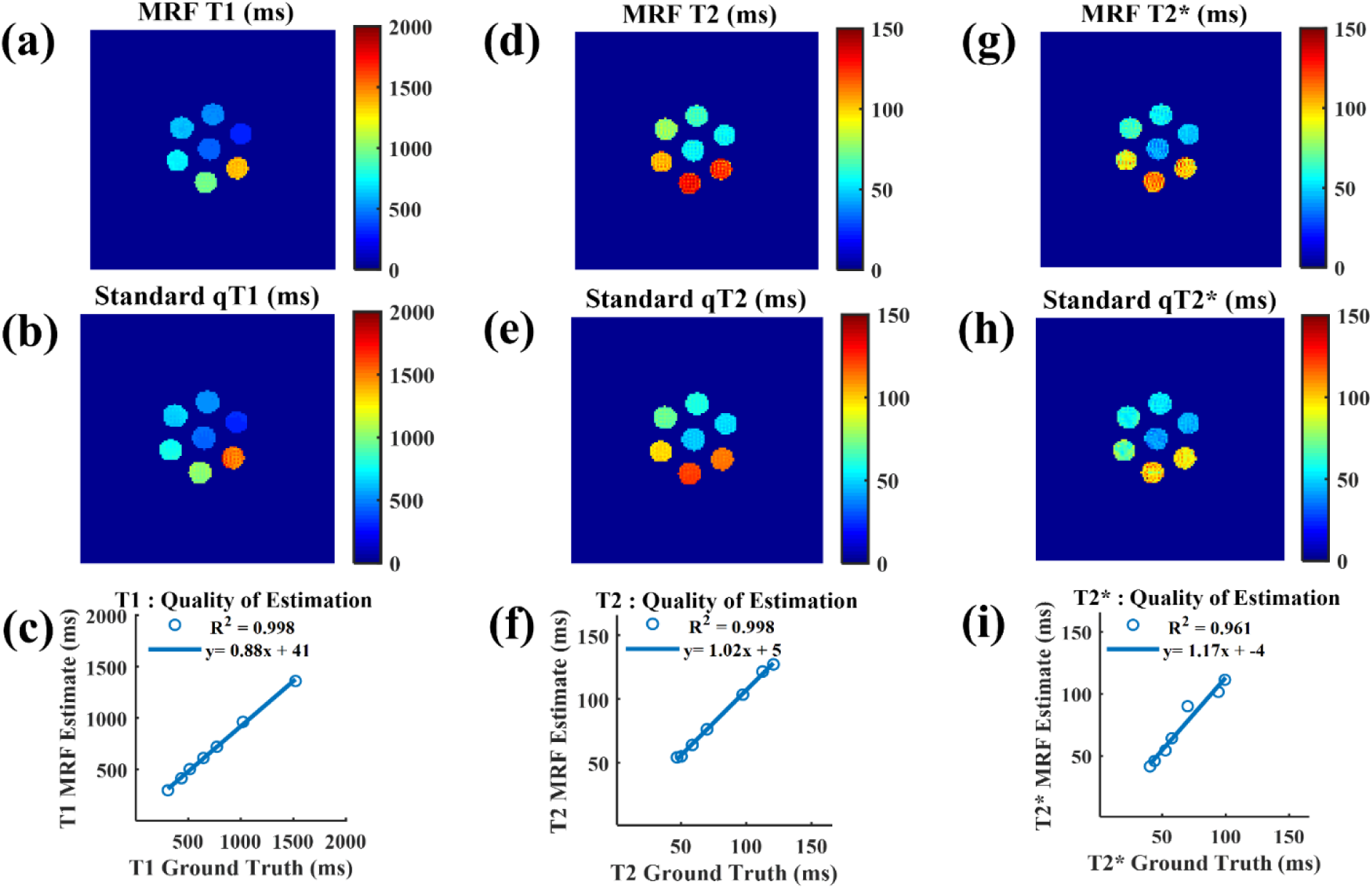
A strong correlation between ground truth approaches to measure T1, T2 and T2* and the estimates of our method was found in a phantom study. The first row shows the spatial parametric maps from our proposed method. The second row shows the results of the ground truth method. The third row shows the correspondence between these two for T1, T2, and T2* measurements, separately.

**Figure 4.**
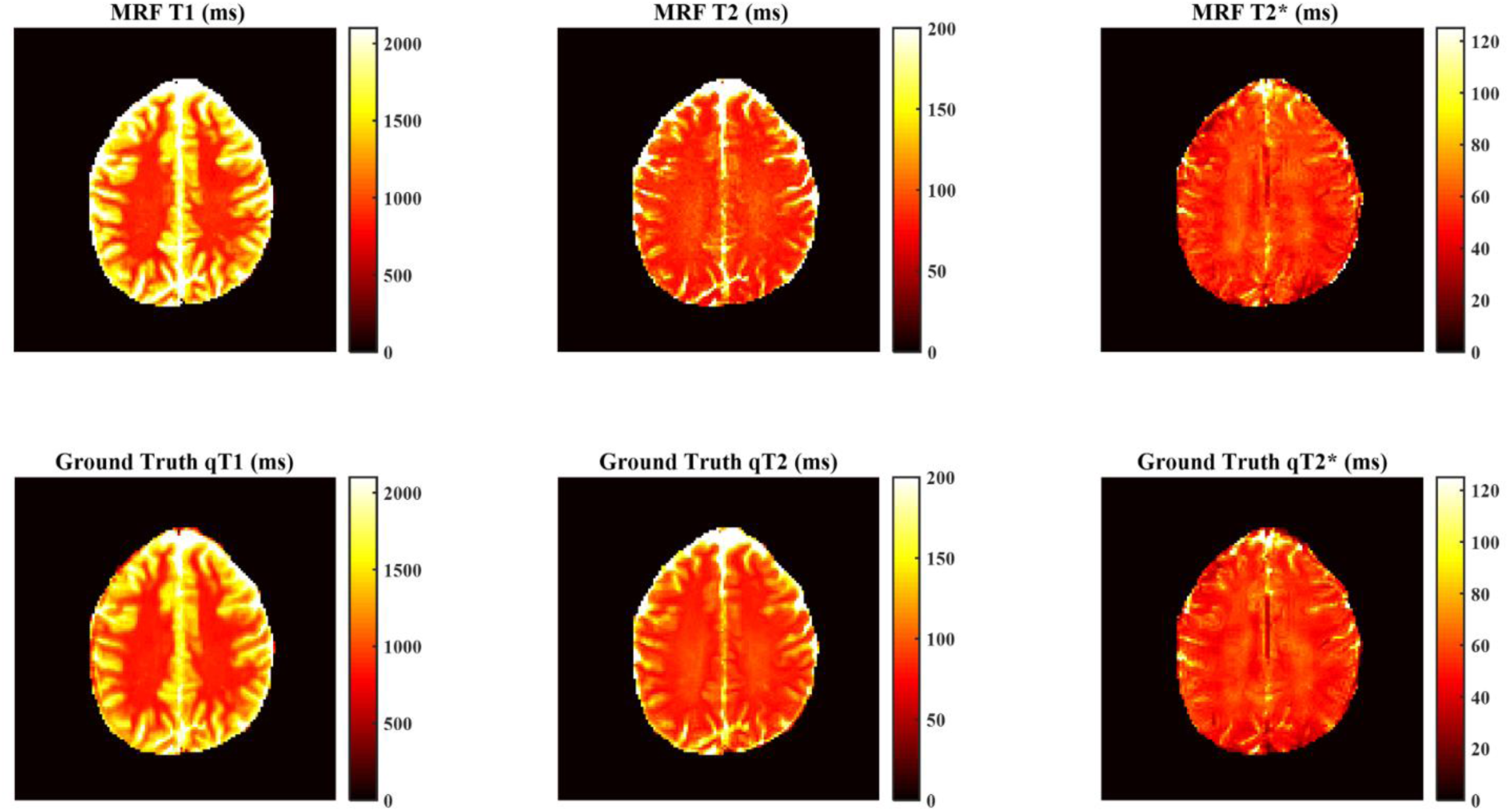
Estimated parametric maps from our proposed MRF method on a human subject (first row) as compared to ground truth measurements for T1, T2 and T2* (second row). Strongly similar contrast and the range of values can be seen between our approach and the ground truth measurements.

Table 1 shows the mean and standard deviation of MRF estimates for GM and WM ROIs as well as the ground truth (relaxometry) values in those regions. No significant differences were found between the two approaches.

**Table 1.** The correspondence between the estimates of our proposed MRF method with the ground truth approaches in a GM- and WM-ROI.

|  | **T1 (ms)** | **T2 (ms)** | **T2\* (ms)** |
| --- | --- | --- | --- |
| MRF WM | $947 \pm 125$ | $79 \pm 4$ | $55 \pm 3$ |
| Ground Truth WM | $913 \pm 108$ | $81 \pm 3$ | $53 \pm 5$ |
| MRF GM | $1153 \pm 154$ | $88 \pm 7$ | $56 \pm 4$ |
| Ground Truth GM | $1122 \pm 52$ | $89 \pm 11$ | $53 \pm 3$ |

## 4) Discussion

In this work, by using an integrated GE-SE sequence, we extend EPI-based MRF to incorporate T2 estimation in addition to that of T1 and T2*. While the advantages of spiral imaging are well known and discussed earlier, there are several arguments in favor of using EPI readout for MRF as compared to undersampled spirals (18,24). As discussed previously, EPI readout is readily available on commercial systems with manufacturer-provided approaches for correcting gradient delays, imperfections and nonlinearities. In addition, due to the absence of undersampling artifacts, far fewer imaging volumes were found to be necessary for accurate parameter estimation (18,25). This leads to faster dictionary generation, lower storage requirements and faster dictionary matching. For example as a rough comparison, in our case, the dictionary size is over three orders of magnitude smaller than a previously reported study (16) that used spirals to estimate the same tissue relaxation parameters (T1, T2, T2*).

Our results show that the acquired signal in EPI-based MRF has high similarity with the dictionary matched entries (see e.g. Figure 2). Owing to the absence of undersampling artifacts, the high image quality of our approach lends itself to the use of accelerated dictionary-matching (26) and more accurate partial volume estimation (27,28).

### 4.1) Limitations and Future Work

While we showed the feasibility of using EPI MRF to estimate T2 in addition to T1 and T2*, we are mindful of the following limitations.

To estimate T2 with EPI readouts, the reversible part of T2* decay should be minimized. We see two different ways to address this. While GE-EPI with a very short TE (e.g. ∼13 ms) can be used along with an optimized pattern of TR/FA change (25,26), off-resonance effects cannot be fully ignored with this approach, and TE minimization is challenged by hardware limitations and the need to ensure uniqueness of dictionary elements. In our approach, a more pure T2 contrast can be achieved using an additional refocusing pulse (i.e. SE-EPI) to compensate for the off-resonance effects. With the TR range involved, this could lead to saturation of the longitudinal magnetization. In our study, we added a wait time (300 ms) after each readout to let the signal recover. Increasing this recovery time could increase the baseline signal, allowing parameter estimation with fewer image volumes. Supplementary Figure 1 demonstrates this effect for three different recovery times (0, 300ms and 600ms) using Bloch simulations. The increased baseline signal due to the increase in the recovery time can clearly be seen in this figure. In addition, this wait time could be used to acquire more slices (19).

In our study, the pattern of TR/TE/FA variations was not optimized. Optimizing it could make the approach faster and more efficient without sacrificing accuracy. In this case, several aspects need to be accounted for, including baseline signal level (which affects image and temporal-SNR), separability of the dictionary elements based on temporal and spatial noise distributions, hardware limitations, slice profile imperfections or magnetization transfer effects. Therefore, finding a globally optimized pattern for EPI MRF (or MRF in general) will not be trivial but will be worth pursuing in future research.

## 5) Conclusion

In this work, we presented a combined GE-SE MRF pulse sequence to estimate T2 in addition to T1 and T2* using an EPI-based MRF framework. This study shows that by providing a robust estimate of all tissue parameters of interest (i.e., T1, T2 and T2*), EPI-MRF can be a viable alternative to spiral based MRF.

## Acknowledgments

We are grateful for financial support from the Canadian Institutes of Health Research (JJC), the Sandra Rotman Foundation and the University of Toronto (MK).

