## Supplementary Figure 1 for "Magnetic Resonance Fingerprinting with Combined Gradient- and Spin-echo Echo-planar Imaging: Simultaneous Estimation of T1, T2 and T2* with integrated-B1 Correction"

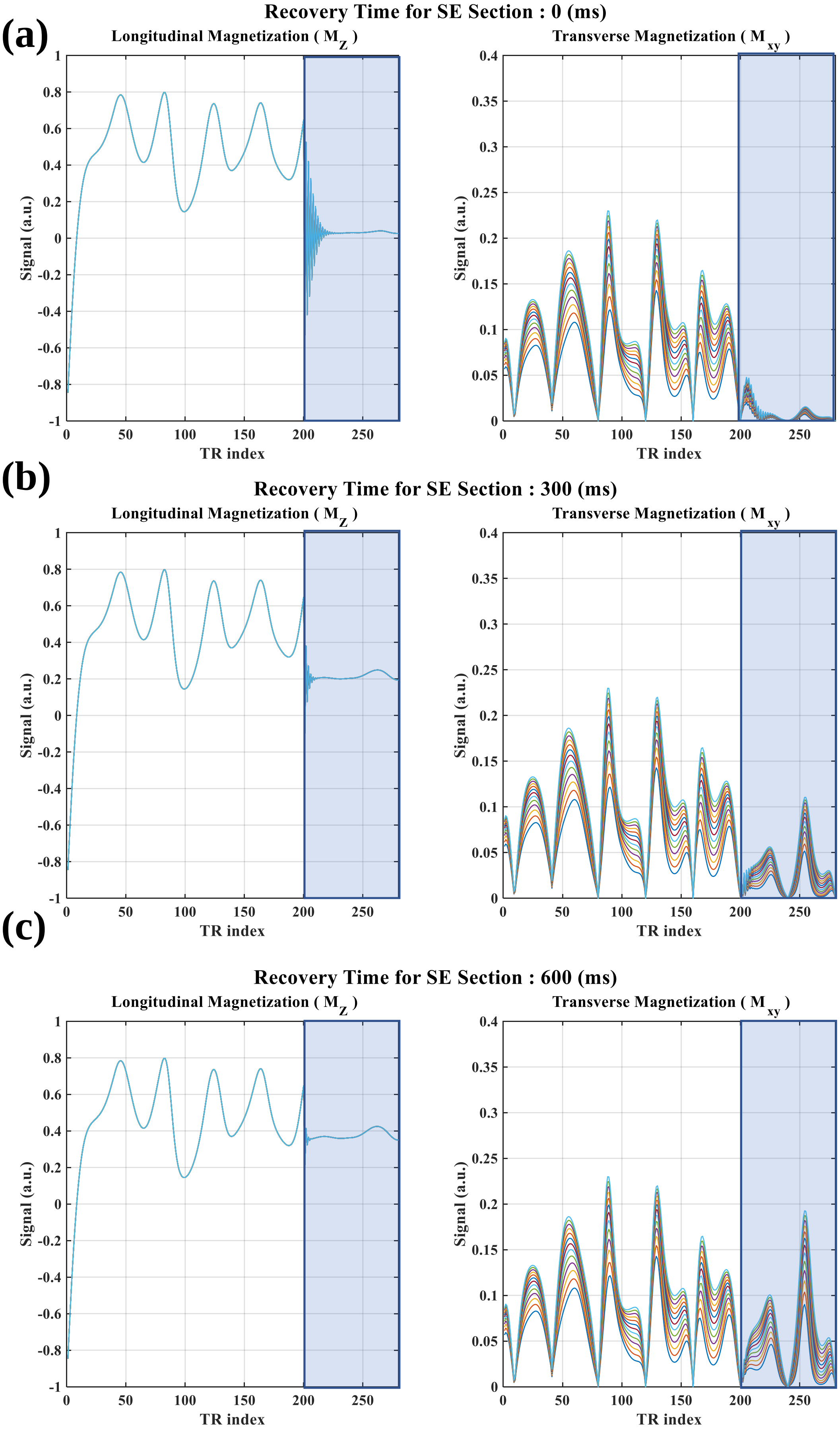


**Supplementary Figure 1**. The relationship between the longitudinal and transverse magnetization and the recovery time when T1=950 ms and T2 varies between 40-100 ms, explored through Bloch simulations. Recovery time = 0 ms (a). Recovery time = 300 ms (b). Recovery time=600 ms(c). Shaded region shows SE section. Increased baseline signal in the SE part due to increased recovery time can be clearly noticed here.
